# Generation of SIV resistant T cells and Macrophages from Nonhuman Primate Induced Pluripotent Stem Cells with Edited CCR5 locus

**DOI:** 10.1101/2021.05.03.442446

**Authors:** Saritha S. D’Souza, Akhilesh Kumar, Jason Weinfurter, Mi Ae Park, John Maufort, Lihong Tao, HyunJun Kang, Thaddeus Golos, James A Thomson, Matthew Reynolds, Igor Slukvin

## Abstract

Adoptive therapies with genetically modified somatic T cells rendered HIV resistant have shown promise for AIDS therapy. A renewable source of HIV resistant human T cells from induced pluripotent stem cells (iPSCs) would further facilitate and broaden the applicability of these therapies. Here, we report successful targeting of the CCR5 locus in iPSCs generated from peripheral blood T cells (T-iPSCs) or fibroblasts (fib-iPSCs) from Mauritian Cynomolgus macaques (MCM), using CRISPR/Cas9 technology. We found that CCR5 editing does not affect pluripotency or hematopoietic and T cell differentiation potentials of fib-iPSCs. However, deletion of CCR5 in T-iPSCs leads to selective loss of their T cell redifferentiation potential without affecting myeloid development. T cells and macrophages produced from CCR5-edited MCM- iPSCs did not support replication of the CCR5-tropic simian immunodeficiency viruses SIVmac239 (T-cell tropic) and SIVmac316 (macrophage-tropic). Overall, these studies provide a platform for further exploration of AIDS therapies based on gene-edited iPSCs in a nonhuman primate preclinical model.

## Introduction

Adoptive T cell therapies with *in vitro* expanded genetically-modified T cells have been considered a valuable strategy to treat and cure HIV.^1-4^ However, T cell exhaustion along with complicated logistics for generation and delivery of genetically modified T cells hampers the broader application of these technologies. Genetic modification of induced pluripotent stem cells (iPSCs) to introduce HIV-resistance and/or anti-HIV molecules, can serve as a versatile and scalable source for off-the-shelf adoptive T cells therapies. In addition, reprogramming of HIV-specific cytotoxic T lymphocytes (CTLs) from HIV-infected patients allows for capturing the specific TCRs within the iPSC genome and generating “rejuvenated” antigen-specific CTLs from these iPSCs.^5, 6^

To enable evaluation of iPSC-based technologies in a preclinical HIV infection model, we explored the feasibility of interrupting the CCR5 locus in iPSCs from nonhuman primate (NHP) sources and *de novo* generating simian immunodeficiency (SIV) resistant T cells and macrophages from these modified iPSCs. In these studies, we used Mauritian cynomolgus macaques (MCM) which have a limited major histocompatibility complex (MHC) diversity,^7-9^ and could be used to assess adoptive cellular therapies, including T cell therapies in a MHC defined setting.^10^ iPSCs were generated from fibroblasts and peripheral blood T cells. To successfully disrupt *CCR5*, we designed two CCR5 synthetic guide RNAs (gRNAs) to target sequences within exon 2, including a 24-bp deletion region that was previously found to prevent functional CCR5 expression in NHPs.^11^ Using this approach, we generated SIV resistant T cells and macrophages from NHP-iPSCs, thus laying a foundation for further exploration of iPSC technology for AIDS treatment in NHP preclinical model. In addition, we noted an impaired capacity of iPSCs generated from T cells (T-iPSCs) to re-differentiate into T cells, especially following biallelic CCR5-disruption. This finding should be taken into consideration when designing strategies for HIV immunotherapies using rejuvenated T cells.

## Results

### Editing of *CCR5* locus in iPSCs from MCM fibroblasts and T cells

Using Sendai virus and oriP/EBNA-1 episomal plasmids, we successfully established iPSC lines from both fibroblasts and peripheral blood T cells from an MCM with an M3/M3 genotype (fib-iPSC-M3/M3 and T-iPSC-M3/M3) and from T cells from an MCM with MHC M1/M3 genotype (T-iPSC-M1/M3) (Figure 1A, Supplemental Figure 1). Generated fib-iPSCs and T-iPSCs exhibited typical NHP embryonic stem cell (ESC) morphology and expressed pluripotency markers OCT4, SOX2 and NANOG (Supplemental Figure 2A-C).

**Figure 1.**
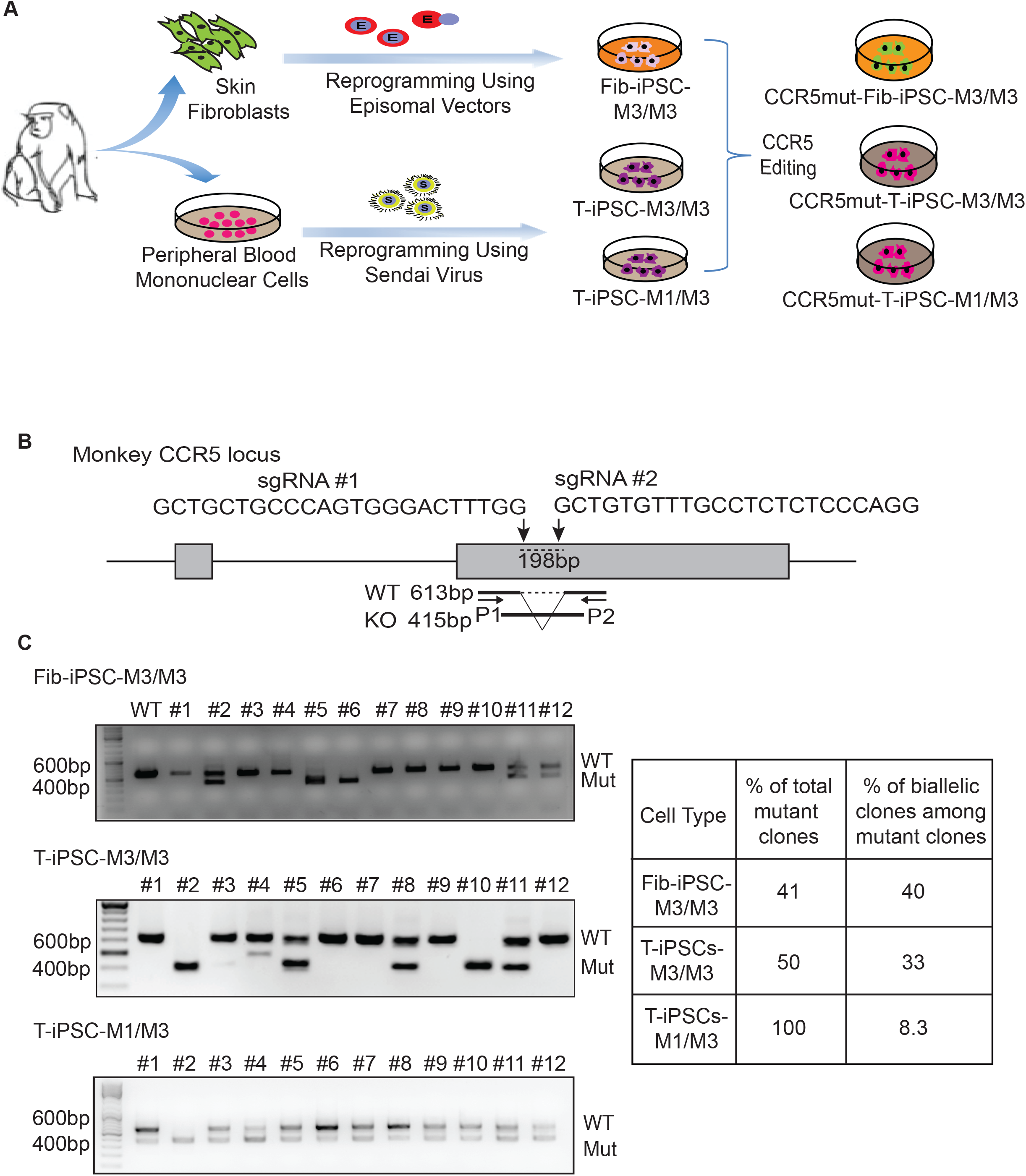
Generation of CCR5mut iPSCs from MCM fibroblasts and T cells. (A) Schematic diagram of experimental design. (B) Schematic representation showing the target site, sequences of the two gRNAs used to delete 198bp CCR5 fragment, and position of PCR primers (P1 and P2) used to detect deletion. (C) Genomic PCR to detect deletion within CCR5 locus. 12 clones from each iPSC line were analyzed.

To disrupt CCR5 we used two gRNAs to target sequences within exon 2, including a 24-bp deletion region known to be essential for expressing functional CCR5 in NHPs ^11^ (Figure 1B). We have shown in prior studies with human iPSCs that dual sgRNAs were more efficient in introducing CCR5 gene editing in human iPSCs as compared to a single sgRNA.^12^ By genomic PCR, we found that the dual gRNAs resulted in 41% of CCR5 mutations in fib-iPSCs-M3/M3, 50% in T-iPSC-M3/M3, and 100% in T-iPSCs M1/M3. Of these, 40% of the clones demonstrated biallelic mutation in the Fib-iPSC-M3/M3 and 33% in T-iPSC M3/M3 and 8.3% in T-iPSC M1/M3 (Figure 1C). Following CCR5 editing, iPSCs retained pluripotent morphology and expression of pluripotency markers OCT4, SOX2 and NANOG (Supplemental Figure 2A and C). Karyotyping revealed a normal karyotype for CCR5mut-fib-iPSC-M3/M3 and CCR5mut-T-iPSC-M1/M3. However, CCR5mut-T-iPSC-M3/M3 demonstrated a balanced translocation between the long (q) arms of chromosomes 2 and 7 (Supplemental Figure 2D). This translocation was detected in several CCR5 mutated clones and was traced back to the original T-iPSC-M3/M3 line suggesting that it was introduced during reprogramming, rather than during CCR5 editing.

### Generation of T cells and macrophages from CCR5-knockout iPSCs

To induce hematopoietic differentiation, we used the OP9 coculture system with CHIR99021 and VEGF to efficiently induce mesoderm and definitive hematopoiesis^13^ (Figure 2A). Floating cells collected from day 10 of iPSCs/OP9 coculture were analyzed by flow cytometry. All iPSCs, including wild-type and CCR5mut fib- and T-iPSCs efficiently produced multipotent hematopoietic progenitors (MHPs) with more than 90% of floating cells expressing CD34 and CD45 (Figure 2B).

**Figure 2.**
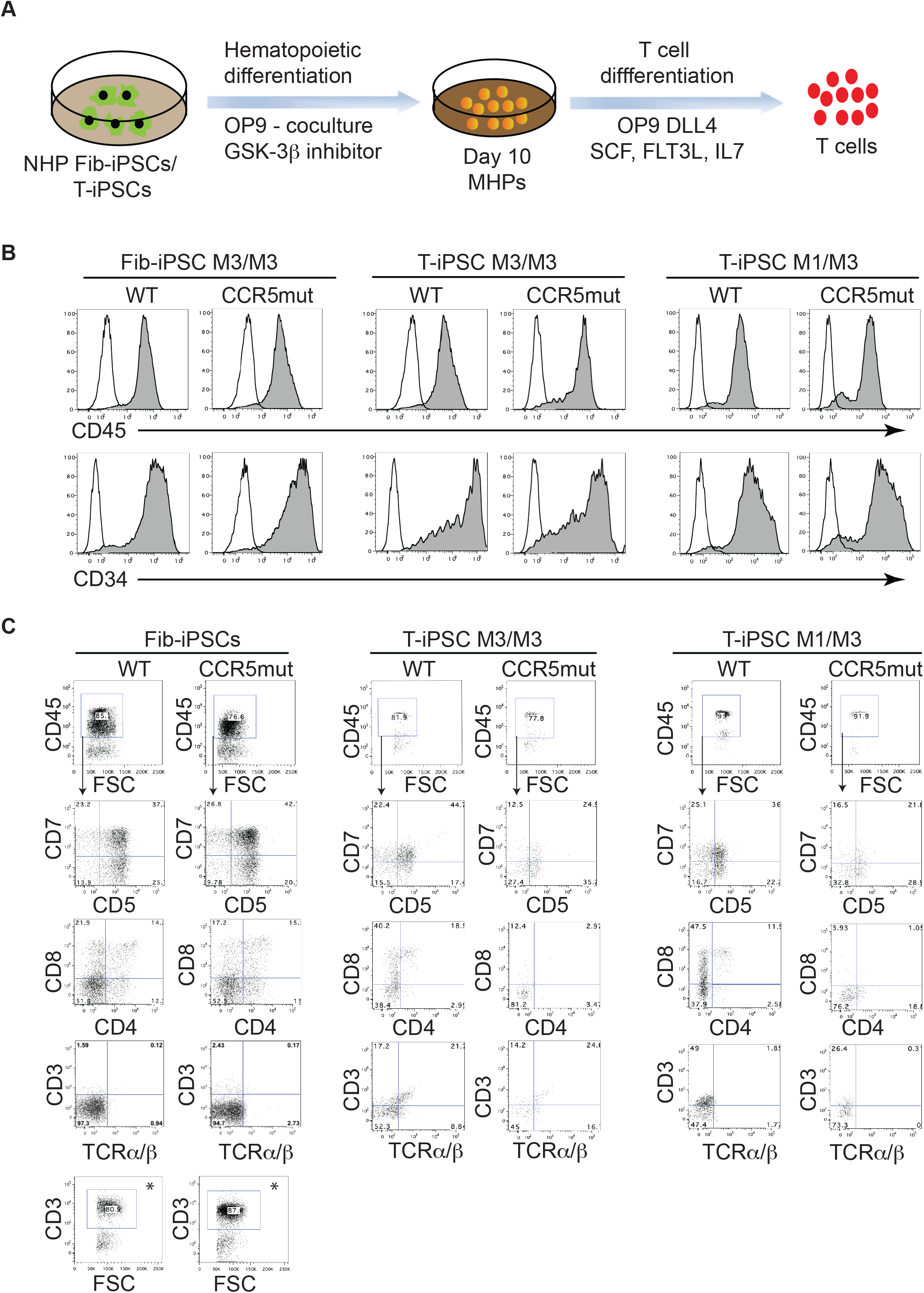
Generation of T cells from CCR5mut iPSCs. (A) Schematic representation of hematopoietic differentiation of fib- and T-iPSCs to MHPs and their further differentiation to T cells. (B) Both WT and CCR5mut iPSCs efficiently differentiated into MHPs as analyzed by flow cytometry of floating cells collected from iPSC/OP9 cocultures on day 10 of differentiation. (C) Day10 MHPs from WT and CCR5mut iPSCs were cultured on OP9-DLL4 in the presence of SCF, IL7 and FLT3L for 2 weeks and the floating cells were analyzed by flow cytometry after gating CD45^+^ cells. * Intracellular CD3 was analyzed in WT and CCR5mut fib-iPSCs. In (B) and (C) Control staining with the appropriate isotype matched mouse monoclonal antibody were included to establish a threshold for positive staining. The graphs are representative of at least 3 independent experiments.

To induce T cell differentiation, we collected day 10 CD34^+^CD45^+^ MHPs and cultured them on OP9-DLL4 in the presence of IL7, FLT3 ligand and SCF according to our protocol which generates functional T cells with rearranged TCR.^13, 14^ In these cultures, MHPs gave rise to CD5^+^CD7^+^ lymphoid progenitors and eventually to CD4^+^CD8^+^ T cells (Figure 2C). As reported in prior studies with human T-iPSCs,^5^ MCM T-iPSCs show surface CD3 expression very early during differentiation. As shown in Figure 2C, surface CD3 expression was already detected in T-iPSC cultures at 2 weeks of T cell differentiation. However, T cells from fib-iPSCs demonstrated mostly intracellular CD3 expression with negligible surface CD3 expression at a similar stage of differentiation (Figure 2C). No differences in T cell differentiation were observed between wild-type fib-iPSC-M3/M3 and CCR5mut-fib-iPSC-M3/M3. However, T cell differentiation of MHPs from T-iPSCs was less efficient as compared to fib-iPSCs and both CCR5mut-T-iPSC-M1/M3 and CCR5mut-T-iPSC-M3/M3 iPSCs failed to produce CD4^+^CD8^+^ T cells (Figure 2C).

For macrophage differentiation, the day 10 MHPs were collected and plated on ultralow attachment plates in IMDM with 10% FBS, M-CSF and IL1β (Figure 3A. Cells collected after 5-7 days displayed typical macrophage morphology and phenotype (Figure 3B). We did not observe significant differences in the macrophage differentiation between wild-type and CCR5mut-fib-iPSCs and T-iPSCs. Genomic PCR analysis confirmed that biallelic CCR5 mutations were maintained in the both the fib-iPSC and T-iPSC derivatives following hematopoietic differentiation including, multipotent CD34^+^CD45^+^ MHPs, macrophages and T cells (Figure 3C and 3D). In addition, the presence of native CCR5 mRNA in macrophages and T cells from wild- type iPSCs and the lack of CCR5 mRNA expression in these cells from CCR5-mut iPSCs was confirmed by RT-PCR (Figure 3E).

**Figure 3.**
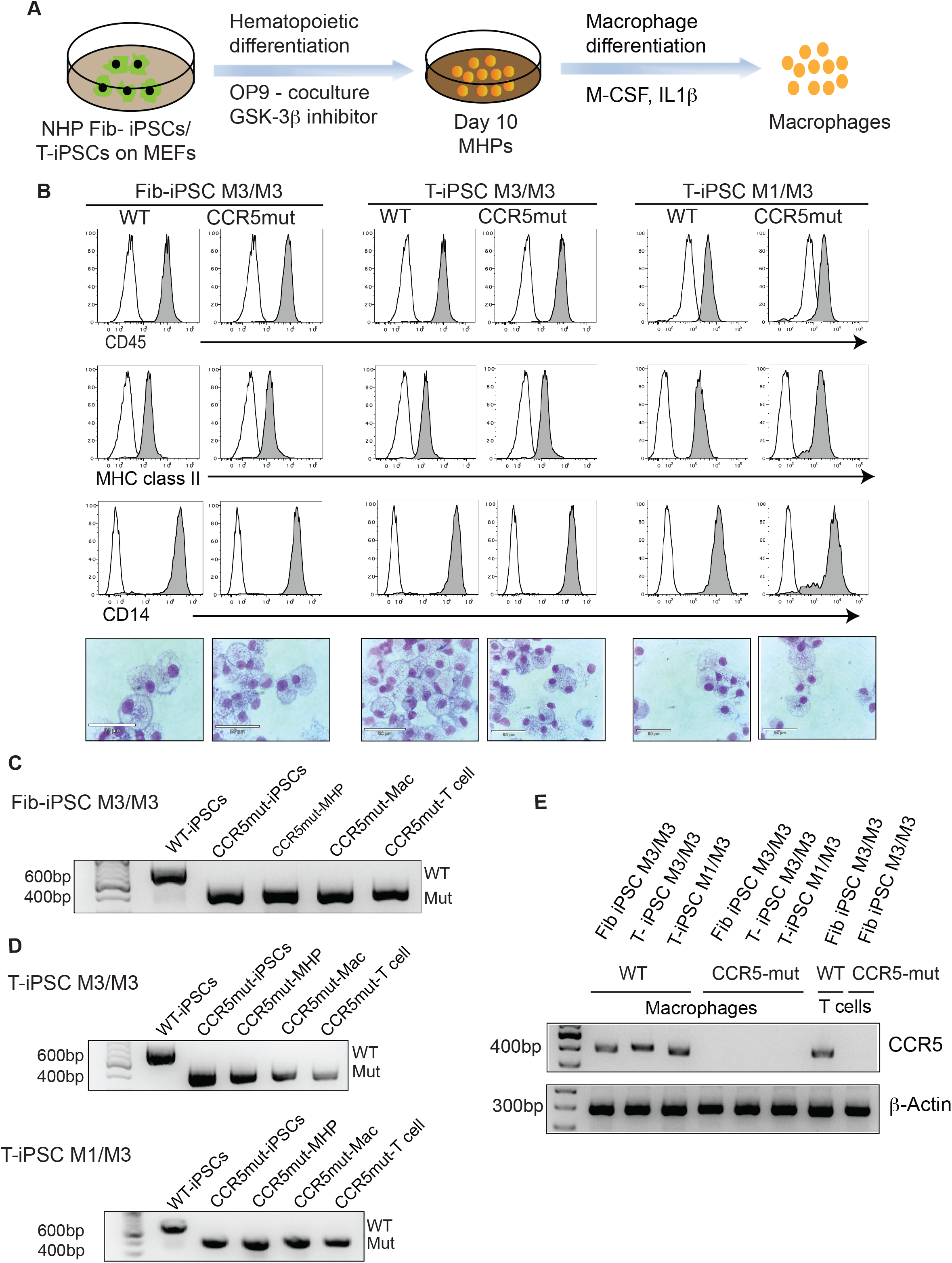
Generation of macrophages from CCR5mut iPSCs (A) Schematic representation of hematopoietic differentiation of iPSCs to MHPs and their further differentiation into macrophages. (B) Day10 MHPs were cultured for 5-6 days in the presence of M-CSF and IL1β to generate macrophages. The phenotype and morphology of the cells was confirmed by flow cytometry and Wright Stain. Representative graphs and images of 3 independent experiments are shown. (C&D) Biallelic CCR5 mutation was maintained in the iPSCs, MHPs, macrophages and T cells following differentiation. (E) Loss of CCR5 expression was confirmed in macrophages and T cells from both wild type and mutant clones by RT-PCR. β-actin was used as an internal control.

### Resistance of CCR5 knockout macrophages and T cells to SIV infection

The aim of our study was to generate CCR5-gene disrupted NHP-iPSCs that when differentiated to T cells and macrophages are resistant to CCR5-tropic SIV infection. Since CCR5mut-T-iPSCs failed to differentiate into T cells, we used only fib-iPSC-M3/M3 for these studies. To evaluate protection from infection, we challenged T cells and macrophages from wild-type and CCR5-mutated fib-iPSCs with the T cell-tropic SIVmac239 and macrophage-tropic SIVmac316 open SpX (1) virus isolates. As shown in Figure 4, SIV replication, as judged by SIV Gag p27 production, was observed at 4 days in T cells and macrophage cultures from wild-type fib-iPSCs. In contrast, CCR5-mutated T cell and macrophage cultures were resistant to productive SIV infection (Figure 4). Although we detected very low levels of p27 in CCR5-mut cell cultures, we believe that this is due to residual virus remaining bound to the outside of cells after magnetofection with high virus concentration. However, it is theoretically possible that a low level of virus replication is still occurring in CCR5-mut immune cells. Overall, these results demonstrate that *CCR5* gene-disrupted T cells and macrophages successfully generated from CCR5 gene altered MCM-iPSCs were protected from CCR5-tropic SIV challenge.

**Figure 4.**
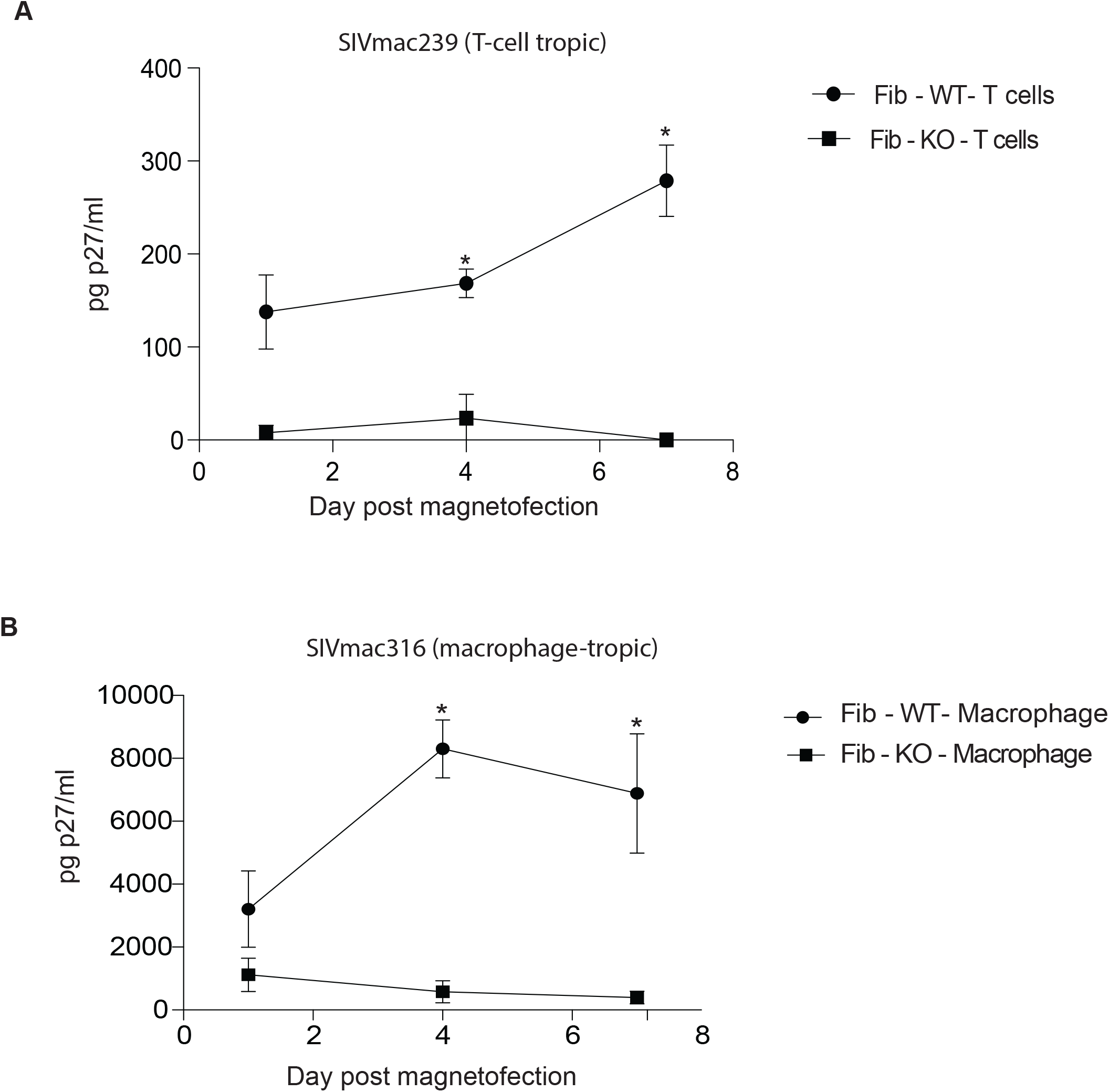
CCR5 mutant T cells and macrophages resist SIV infection. Fib-iPSC-derived mutant and wild-type T cells (A) and macrophages (B) were incubated with T-cell-tropic SIVmac239 and macrophage-tropic SIVmac316, respectively. Virus production was measured by collecting cell culture supernatant on days 1, 4, and 7 post-infection and performing SIV Gag p27 ELISAs. The assay was performed in triplicate and the error bars represent the standard error of each time point. * denotes P<0.05.

## Discussion

iPSCs derived from somatic cells offers an attractive strategy for the generation of HIV-resistant immune cells for adoptive immunotherapies. In this study, we demonstrated the feasibility of reprogramming T cells from NHPs to pluripotency and generation of SIV resistant CCR5-mutated T cells and macrophages from fib-iPSCs obtained from MCMs. MCMs are descended from a small founder population and have a very limited MHC diversity consisting of only seven common haplotypes M1-M7,^7, 8^ therefore making it possible to rapidly select MHC identical, MHC homozygous and MHC heterozygous animals to explore the utility of MHC homozygous iPSC banking for allogeneic immunotherapies using cells with beneficial MHC match in a preclinical NHP model.

Gene editing to inactivate the CCR5 gene using nucleases has shown promising results towards developing a functional HIV cure.^15-18^ Three main classes of nucleases, including ZFNs, TALENs and CRISPR/Cas9 are routinely used to precisely delete or insert specific DNA sequences into genome. Of these, nucleases, the CRISPR/Cas9 system has demonstrated higher cleavage efficiency compared to ZFNs and TALENs.^19-22^ In this study, we employed dual gRNA- guided Cas9 systems to specifically target the CCR5 region affected by Δ24 mutation, which prevents R5 lentiviral infections in macaques^11^ and is functionally equivalent to the *CCR5*Δ*32* mutation in humans.^23, 24^ Using the dual gRNA CRISPR/Cas9 system we achieved up to a 40% biallelic mutation in NHP iPSCs. Our results are consistent with several other reports showing improved editing efficiency using dual gRNA in primary human CD4^+^ T cells, CD34^+^ hematopoietic progenitor cells, and iPSCs^12, 25, 26^ T cells and macrophages produced from CCR5mut-fib-iPSCs were resistant to SIV challenge, implying that immune cells derived from the NHP-iPSCs are useful for continued studies of novel immunotherapies for HIV in a preclinical NHP model.

Generating T cells from T-iPSCs opens opportunities for T cell rejuvenation and unlimited manufacturing of T cells with an antigen-specific monoclonal TCR.^5, 27, 28^ The feasibility of this approach for generating therapeutic cells has been demonstrated in murine and human studies, where rejuvenated T cells from tumor- or virus-specific targets *in vivo* and *in vitro,*^27, 29, 30^ including HIV.^5^ To advance this strategy in a preclinical NHP model, we successfully generated T-iPSCs from MCM peripheral blood T cells, and demonstrated that T-iPSC can be successfully re-differentiated into T cells. However, we found that MHP generated from T-iPSC produced T cells less efficiently and CCR5mut-T-iPSCs did not generate T cells, despite no apparent loss in the efficiency of hematopoietic differentiation and macrophage production from wild-type and CCR5mut-T-iPSCs. This defect in hematopoietic differentiation affected the T cell differentiation step only from T-iPSCs, but not fib-iPSCs, which produced T cells efficiently regardless of the presence or absence of the CCR5 mutation. The reason for these differences remains unclear. It has been shown that following differentiation, T-iPSCs express TCR complex prematurely before the CD4^+^CD8^+^ double-positive stage, ^29, 31^ which may lead to strong TCR-signaling and eventually death of T cell progenitors. In addition, previous studies have revealed that CCR5 expression promotes IL2-dependent events during T cell activation, including IL2R expression, STAT5 phosphorylation and T cell proliferation.^32^ Thus, it is possible that CCR5 deficiency in an environment of premature TCR expression may further contribute to the demise of T cell differentiation potential.

In summary, we have shown that genomic editing of *CCR5* can be easily and effectively attained in NHP iPSCs which can be clonally selected to ensure homogenous CRISPR/Cas9 gene editing. These lines can be useful for understanding the role of CCR5 in HIV pathogenesis and further advancement of iPSC-based technologies for a HIV cure in NHP preclinical models. We also noted that introduction of CCR5 mutation into T-iPSCs affects their T cell redifferentiation potential. This unexpected finding presents an additional challenge to applying T cell rejuvenation technologies for HIV therapies using CCR5-edited HIV-resistant T-iPSCs.

## Material and Methods

### Reprogramming NHP cells and demonstration of pluripotency

Fibroblasts were obtained from MCM CY1 with M3/M3 MHC genotype, while peripheral blood samples were obtained from CY1 and CY2 with corresponding M3/M3 and M1/M3 MHC genotypes. Fibroblasts were obtained from skin biopsy and reprogrammed using a combination of oriP/EBNA-1 episomal vectors as described previously^13, 33^ to generate fibroblast iPSCs (Fib-iPSC). Peripheral blood T cells were reprogrammed using the CytoTune™-iPS 2.0 Sendai Reprogramming Kit (ThermoFisher Scientific) to generate T-iPSCs.^34^ Briefly, peripheral blood mononuclear cells were first separated by Ficoll centrifugation and then MACS enriched for CD3^+^ cells. These cells were then activated using CD3 and CD28 antibodies in the presence of 100U/ml IL-2 (Peprotech). For reprogramming, 0.5×10^6^ activated cells were resuspended in 0.3ml of 10% RPMI1640 medium (ThermoFisher) containing 4μg/ml of hexadimethrine bromide (MilliporeSigma) and 20μl each of Sendai virus KOS (KLF4–OCT3/4–SOX2), cMYC and KLF4, and incubated overnight at 37°C in 5% CO_2_. Cells were washed the following day and transferred onto mouse embryonic fibroblasts (MEFs). Colonies began to appear within 15 days and were subsequently transferred onto fresh MEFs for expansion and characterization. The iPSC lines (T-iPSC and Fib-iPSC) were maintained on MEFs in Primate ES cell medium (ReproCELL) supplemented with 4ng/ml bFGF (154 a.a.) (Peprotech) as previously described.^13^ Cells were passaged every 3-4 days using Collagenase Type IV (Life Technologies). Expression of pluripotency markers was analyzed by flow cytometry using SOX2 (Cell Signaling) and OCT3/4 (Santacruz Biotechnology) antibodies and the MACSQuant Analyzer 10 (Miltenyi) and FlowJo software (BD). Expression of pluripotency marker NANOG was analyzed by immunofluorescence using antibodies from Cell Signaling.

### NHP iPSC culture and generation of CCR5 CRISPR knockout iPSC line

NHP iPSCs were harvested using Collagenase IV followed by TrypLE to make a single cell suspension. 1×10^5^ singularized cells were resuspended in 100μl of nucleofector solution (Lonza) containing 10μg of each sgRNA (#1 and #2 modified sgRNAs, Synthego) and 15μg Cas9 protein (PNA Bio) and were electroporated using program A23 on the Nucleofector™ 2b Device (Lonza). After transfection, cells were replated onto MEFs in Primate ES cell medium supplemented with 4 ng/ml bFGF (154 a.a.). 15-20 days later, colonies were picked and expanded. Single cell derived KO cell lines were obtained by single colony picking method with low density iPSCs culture on MEFs. Genomic DNA from iPSC colonies was extracted using the Quick-DNA Miniprep kit (Zymo Research) and analyzed by PCR. The targeting genomic PCR in CCR5-mutated clones was performed using Q5 Hot Start High Fidelity DNA polymerase (NEB) with the following primers: P1 (TCAATGTGAAACAAATCGCAGC) and P2 (TCGTTTCGACACCGAAGCAG).

### Differentiation to T lymphoid cells and Macrophages

Hematopoietic differentiation of NHP iPSCs was performed on OP9 in the presence of CHIR99021 as previously described.^13^ Briefly, small cell aggregates of iPSCs were added to a prolonged culture of OP9 feeder in medium supplemented with 10% HyClone™ FBS (Cytiva) and 50μM β-mercaptoethanol (MilliporeSigma). 4μM of CHIR99021 (Peprotech) and 50ng/ml VEGF (Peprotech) were added on day 1, for 2 days. The medium was changed and fresh medium supplemented with 50ng/ml VEGF was added. On day 6, an additional 5ml of medium along with a hematopoietic cytokine cocktail consisting of 50ng/ml SCF (Peprotech), 50ng/ml of VEGF (Peprotech), 20ng/ml of TPO (Peprotech), 20ng/ml of IL-3 (Peprotech) and 20ng/ml of IL-6 (Peprotech) was added to the co-culture. The co-culture was incubated for 10 days in standard conditions of 37°C and 5% CO_2_. The phenotype of the cells was confirmed by flow cytometry using antibodies against CD45 (Miltenyi) and CD34 (BD Biosciences).

For lymphoid differentiation, the floating CD45^+^CD34^+^ multipotent hematopoietic progenitors (MHPs) were collected from day 10 of NHP iPSC/OP9 cocultures, strained through a 70μm cell strainer (ThermoFisher Scientific) and resuspended in a T cell differentiation medium consisting of αMEM (Gibco) supplemented with 20% HyClone™ FBS, 5ng/ml IL-7 (Peprotech), 5ng/ml Flt3-Ligand (Peprotech) and 10ng/ml SCF (Peprotech). The cells were cultured on OP9-DLL4 for 2 weeks with weekly passage. The floating cells from T cell cultures were analyzed by flow cytometry using antibodies against CD3, CD4, CD7 (BD Biosciences), CD5, CD7 and TCRαβ (Biolegend) and used for subsequent SIV challenge. For intracellular staining, cells were fixed and permeabilized by resuspending the cell pellet in BD cytofix/cytoperm buffer (BD Biosciences) for 30 mins on ice. The cells were then washed with 1×Perm Buffer (BD Biosciences) and stained with CD3 antibody for 30 mins in the dark, washed and analyzed using the MACSQuant Analyzer (Miltenyi Biotec) and FlowJo software (Tree star). Control staining with the appropriate isotype matched mouse monoclonal antibody and unstained controls were included to establish a threshold for positive staining.

For macrophage differentiation, day 10 MHPs from OP9/iPSC co-culture were suspended in Iscove’s Modified Dulbecco’s Medium (IMDM; Gibco) with 10% HyClone™ FBS supplemented with 20ng/ml of M-CSF (Peprotech) and 10ng/ml of IL-1β for 5-7 days. Cell phenotype was confirmed by Wright-Giemsa staining and flow cytometry using antibodies against CD45 (Miltenyi), CD14 (BD Biosciences) and HLA-DR (BD Biosciences). Cells from day 6 of macrophage culture were used for SIV challenge.

### RT-PCR analysis of CCR5 expression

RNA was isolated using the RNeasy mini Kit (Qiagen) from macrophages and T cells generated from wild type and CCR5-mut iPSCs. cDNA was transcribed using the QuantiTect reverse transcription kit (Qiagen) from all the samples and amplified by PCR using Q5 master mix (New England Biolabs) and forward-TGTGTCAATGGAACTCTTGAC and reverse-TCGTTTCGACACCGAAGCAG primers.

### SIV challenge of iPSC derived T cells and macrophages

To determine whether the CCR5 mutant macrophages and T cells were resistant to infection, we challenged them with the CCR5-tropic SIV isolates SIVmac239 and SIVmac316 open SpX, previously shown to be T-cell- or macrophage-tropic, respectfully (Mori et al., 2000). The SIV stocks were purified by overlaying 127 ng (SIVmac316 open SpX) or 87 ng (SIVmac239) of Gag p27 on 100 ul of a 20% sucrose cushion and centrifuging at 21,000g for 1 hour at 4°C. Media and sucrose were removed and the virus pellet was resuspended in 70μl of PBS. Cell infection was performed using magnetofection. Briefly, 30μl of ViroMag R/L beads (OZ Biosciences) were added and incubated at room temperature for 15 minutes.^35^ During incubation 3×10^5^ wild-type or CCR5-mut cells were placed in a single well of a 24-well plate and centrifuged at 530g for 5 minutes. The virus/bead mixture was then added dropwise to the cells and placed on a magnet for 1 hour at 37°C. When the incubation was finished, the cells were pelleted and washed five times with 1 ml PBS. T cells from iPSC were treated with 0.05% or 0.25% trypsin for 2 minutes at 37 °C to remove bound but non internalized virions. Then 1×10^5^ of each virus/cell combination were then placed into three wells of a 48-well plate containing growth media and incubated for 7 days at 37°C, 5% CO2. Culture supernatants were sampled at days 1, 4 and 7 post-magnetofection. A p27 ELISA (Zeptometrix) was performed on each time point according to the manufacturer’s instructions to determine the amount of virus produced in each well.

## Acknowledgement

This work is supported by funds for National Institute of Health (R24 OD021322, R01 HL142665 and P51OD011106). The following reagent was obtained through the NIH AIDS Reagent Program, Division of AIDS, NIAID, NIH: SIV_mac_316 open SpX from Dr. Ronald C. Desrosiers.

## Author Contributions

S.S.D. generated and characterized CCR5mut T-iPSCs and fib-iPSCs, analyzed iPSC hematopoietic differentiation potential, produced macrophages, interpreted experimental data, made figures and wrote manuscript; A.K. iPSCs from T cells, characterized CCR5mut T-iPSCs and generated T cells from iPSCs; J.W. performed SIV infection studies; M.A.P. generated and characterized CCR5mut T-iPSCs; J.M. generated iPSC from fibroblasts. H.J.K. designed CCR5 targeting gRNAs; T.G. and J.A.T. advised on iPSC generation and characterization. M.R. and I.S. developed the concept, led and supervised studies, analyzed and interpreted data and wrote the manuscript.

