## Supplemental data for "Generation of SIV resistant T cells and Macrophages from Nonhuman Primate Induced Pluripotent Stem Cells with Edited CCR5 locus"

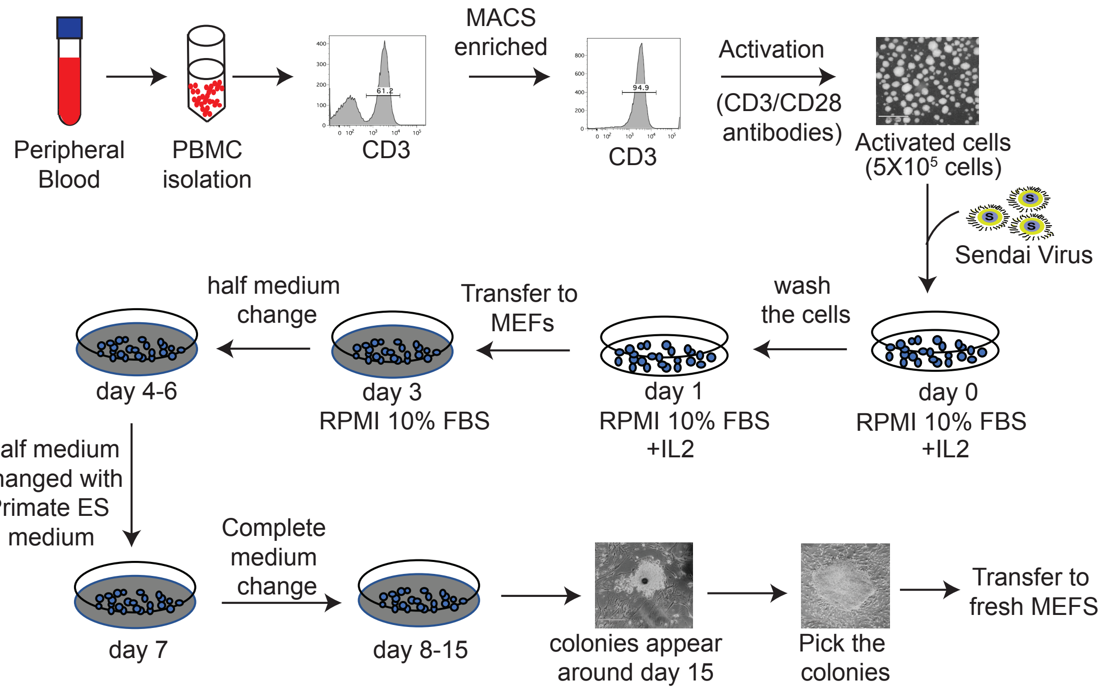


**Supplementary Figure 1.** Schematic diagram of generation of T-iPSCs from peripheral blood T cells using Sendai virus kit.

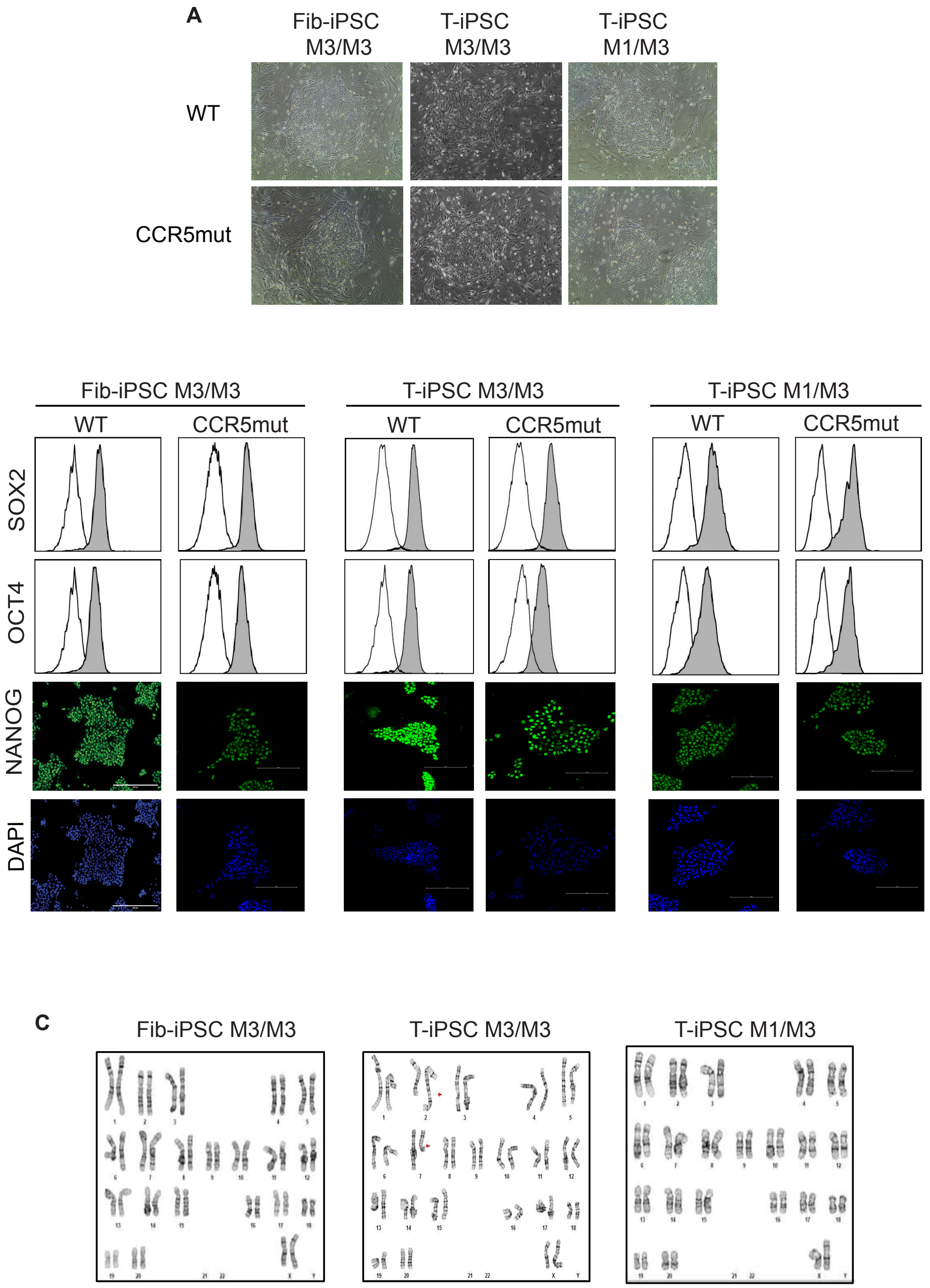


**Supplementary Figure 2.** Generation and characterization of MCM-iPSCs. (A) Phase contrast images of WT and CCR5mut colonies on MEFs. (B) Expression of pluripotency markers by flow cytometry (OCT4 and SOX2) and by immunofluorescence (NANOG) in generated wild type and CCR5mut iPSC lines. (C) Karyotype of CCR5mut iPSCs.
